# Phenology and ecological role of Aerobic Anoxygenic Phototrophs in fresh waters

**DOI:** 10.1101/2023.11.17.567504

**Authors:** Cristian Villena-Alemany, Izabela Mujakić, Livia K. Fecskeová, Jason Woodhouse, Adrià Auladell, Jason Dean, Martina Hanusova, Magdalena Socha, Carlota R. Gazulla, Hans-Joachim Ruscheweyh, Shinichi Sunagawa, Vinicius Kavagutti, Adrian-Ştefan Andrei, Hans-Peter Grossart, Rohit Ghai, Michal Koblížek, Kasia Piwosz

## Abstract

Aerobic anoxygenic phototrophic (AAP) bacteria are heterotrophic bacteria that supply their metabolism with light energy harvested by bacteriochlorophyll-*a*-containing reaction centres. Despite their substantial contribution to bacterial biomass, microbial food webs and carbon cycle, their phenology in freshwater lakes remains unknown. Hence, we investigated seasonal variations of AAP abundance and community composition biweekly across three years in a temperate, meso-oligotrophic freshwater lake. AAP bacteria displayed a clear seasonal trend with a spring maximum following the bloom of phytoplankton and a secondary maximum in autumn. As the AAP bacteria represent a highly diverse assemblage of species, we followed their seasonal succession using the amplicon sequencing of the *puf*M marker gene. To enhance the accuracy of the taxonomic assignment, we developed new *puf*M primers that generate longer amplicons and compiled the currently largest database of *puf*M gene, comprising 3633 reference sequences spanning all phyla known to contain AAP species. With this novel resource we demonstrated recurrent and dynamic seasonal succession of the AAP community. The majority of the species appeared during specific phases of the seasonal cycle, with less than 2% of AAP species detected during the whole year. Our results document the indigenous freshwater nature of the AAP community, characterized by high resilience and heterogenic adaptations to varying conditions of the freshwater environment. By integrating this information with the indicator of primary production (Chlorophyll-*a*) and existing ecological models, we show that AAP bacteria play a pivotal role in the recycling of dissolved organic matter released during spring phytoplankton bloom, contributing significantly to the ecological dynamics of lakes.

## Introduction

Recurrent seasonal changes of aquatic microbial communities are among the best-studied phenomena in freshwater lakes and reservoirs. The Plankton Ecology Group (PEG) model initially described the dynamic interactions between phytoplankton and zooplankton [1] and was later amended with the eutrophic and oligotrophic scenarios and role description of heterotrophic protists [2]. Subsequently, the importance of bacterioplankton was revealed, especially during the spring phytoplankton bloom [3–5], increasing our understanding of the contribution of microorganisms to the functioning of limnic ecosystems [6]. Bacteria represent an important part of aquatic microbial communities. They generate fresh particulate organic matter by utilizing dissolved organic carbon (DOC) and render it accessible to organisms at upper trophic levels [7]. However, the role of photoheterotrophic bacteria, which present a significant part of bacterial biomass and activity [8], remains overlooked.

Aerobic anoxygenic phototrophic (AAP) bacteria are a functional group of photoheterotrophs that rely upon external sources of organic carbon and supplement their metabolism with energy obtained from light through bacteriochlorophyll-*a* (BChl-*a*) type-II reaction centres. This photoheterotrophic life style enables AAP bacteria to reduce their respiration and increase biomass yield in light [9, 10]. Moreover, AAP community show higher growth rates, larger cell sizes and greater activity than heterotrophic bacteria [11–16]. Photoheterotrophy by AAP bacteria increases carbon transfer efficiency, enlarging the availability of biomass for upper trophic levels and reducing CO_2_ emitted to the atmosphere [17]. However, little is known on phenology of AAP community and the absence of exhaustive seasonal sampling hampers the understanding of their role in lakes. AAP bacteria peak during spring in lakes, when they may account for up to 22% of bacteria [18, 19]. Their abundances and diversity dynamics correlate with irradiance, temperature, chlorophyll-*a* (Chl-*a*), oxygen and DOC [18, 20–24].

One of the obstacles in the study of AAP bacteria is the fact that they do not represent a monophyletic group. On the contrary, phototrophic genes have been gained and lost multiple times in closely related species [25, 26]. Therefore, AAP species cannot be identified based on the most common marker used in community studies, 16S rRNA gene. Instead, the *puf*M gene, which encodes the subunit M of the anoxygenic type-II reaction centre, has been widely employed to study AAP communities [15, 27–32]. However, these studies were unsuccessful in providing a taxonomic assignment for abundant *puf*M phylotypes. This is caused by the low taxonomic resolution of the short amplicon sequences and the lack of a curated reference database. The increased availability of metagenome-assembled and single-cell amplified genomes (MAGs & SAGs) has expanded our knowledge of metabolic potential within multiple bacterial lineages and should allow to establish a comprehensive *puf*M database for amplicon assignment.

To improve the taxonomic assignment, we designed a novel primer set targeting a larger 450 bp region of the *puf*M gene and compiled an extensive database of 3633 non-redundant *puf*M gene sequences from existing genome and metagenome sequence datasets. We applied this novel metabarcoding assay to 215 samples from three years, collected from meso-oligotrophic freshwater Cep lake (Czechia) at biweekly intervals from multiple depths. We hypothesized that the AAP community would show a recurrent seasonal succession, marked by distinct phylotypes reaching abundance peaks under specific environmental conditions. Moreover, we expected that AAP bacteria would exhibit different abundance succession patterns than overall heterotrophic bacteria. Specifically, we surmise that the spring AAP bloom is orchestrated by specific phylotypes, rather than the involvement of the entire AAP community.

## Materials and methods

### Sampling and measuring environmental variables

Samples were collected biweekly from April 2017 to December 2019 from the freshwater Cep lake (48°92149.241N, 14°88168.111E). This meso-oligotrophic lake is located in the Třeboň Basin Protected Landscape Area, Czechia, and has an area of 130 ha and a maximum depth of 12 m. Five litres of water were collected from 0.5, 2, 5 and 8 meters using a 3-L Ruttner water sampler (KC Denmark A/S, Denmark) and transported to the laboratory in closed plastic containers in a cooler box, which were pre-rinsed three times with the sampled water. Temperature and oxygen profiles were taken with an EXO1 multi-parameter probe (YSI Inc., Yellow Springs, USA). Total and AAP bacterial abundances were counted using epifluorescence microscopy method as described in Piwosz et al., 2022 [17]. Concentrations of Chl-*a* and BChl-*a* were determined in organic solvent extracts by reversed-phase high-performance liquid chromatography [31]. The quantification of environmental nutrients was performed as described in Procházková, 1959 (nitrate); Murphy and Riley, 1962 (phosphate); Kopáček and Hejzlar, 1993 (total phosphorous); Kopáček and Procházková, 1993 (ammonia) and Shabarova et al., 2021 (DOC) [33–37].

### *puf*M gene database

We collected 14,872 *puf*M nucleotide and protein sequences from representative genomes and MAGs available from Genome Taxonomy Database (GTDB) r207 [38], Tara Ocean [39], the LIMNOS database compiled from a set of ∼1300 freshwater lake metagenomes (PRJEB47226), and from MAG collections publicly available in the NCBI [3, 40–44]. Bacterial genomes and MAGs taxonomy was determined using GTDB-Tk v2.1.1 [45]. *puf*M sequences in GTDB r207 were found using hidden Markov model of *puf*M gene (K08929) from the KOFAM database [46] with a score threshold of 394.57, as described in https://github.com/adriaaula/obtain_gene_GTDB. *puf*M-containing non-redundant MAGs from Tara Oceans (https://doi.org/10.6084/m9.figshare.4902923.v1) were selected using HMMER v3.3.2 (http://hmmer.org/) with a customized *puf*M database from pfam [47]. The *puf*M sequences were confirmed by Diamond v0.9.24 annotation [48]. Nucleotide *puf*M gene sequences from the LIMNOS database were obtained from open reading frames using Prodigal [49] and annotated using a custom pipeline incorporating Diamond v0.9.24 [48] and the KEGG database [50]. *Puf*M genes were compiled alongside the taxonomy of their associated MAG.

All *puf*M sequences were pooled and duplicated sequences were removed. Protein sequences were aligned with MAFFT v7.453 (--maxiterate 1000 --localpair) [51] and a maximum likelihood tree was calculated using iqtree2 [52] with automatic model selection performed by ModelFinder [53], and 1000 iterations of ultrafast bootstrapping with 1000 rounds of SH-aLRT testing (-alrt 1000 −B 1000) [54]. *puf*L sequences were identified as they formed a long branch. Bona-fide *puf*M sequences were retained (Supplementary File S1), and alignment and phylogenetic trees were redone and visualized using iTOL [55]. The environmental origin of each sequence was obtained manually from source databases (Supplementary File S2).

### AAP community analysis by *puf*M gene amplicon sequencing

Between 300 and 1460 ml of water was filtered through sterile 0.2 µm Nucleopore Track-Etch Membrane filters (Whatman®, Maidstone, United Kingdom) that were immediately placed inside sterile cryogenic vials (Biologix Group Limited, Jinan, Shandong China) containing 0.55 g of sterile zirconium beads, flash-frozen in liquid nitrogen and stored at −80 °C until DNA extraction (max. six months). Total nucleic acids were chemically extracted according to Griffiths et al. 2000 [56] with modifications [57], re-suspended in 35 µl of DNase and RNase-free water (MP Biomedicals, Solon, OH, USA) and stored at −20 °C. Concentration and quality of the extracts were checked using NanoDrop (Thermo Fisher Scientific).

To improve the accuracy of the taxonomic assignation and reduce the number of unclassified amplicon sequence variants (ASVs), a new primer pair for *puf*M gene was designed. pufM_uniF primer [27] was used as a reverse (pufM_UniFRC in the current study, 5’-RAANGGRTTRTARWANARRTTNCC-3’) and pufM_longF was designed ∼450 bp upstream (5’-YGGSCCGWTCTAYSTSGG-3’) using a pre-existing database of 1500 sequences [31]. The specificity and coverage of the new primer pair were tested in comparison to the commonly used *puf*M primers [27, 32] against the new *puf*M database. The analysis was done in Geneious (v2023.0.1) with up to three mismatches in the binding region and in both forward and reverse directions. The primers’ specificity was also tested separately for Pseudomonadota (formerly known as Proteobacteria), Alphaproteobacteria, Gammaproteobacteria, Gemmatimonadota, Chloroflexota, Myxococcota and Eremiobacterota based on alignments done in Geneious by MUSCLE alignment (v5.1.).

The PCR conditions were optimized using genomic DNA from *Gemmatimonas phototrophica* (Gemmatimonadota), *Sphingomonas glacialis* (Alphaproteobacteria) and *Congregibacter litoralis* (Gammaproteobacteria), and environmental DNA from the current sampling. The final conditions were as follows: initial denaturation for 3 min at 98°C, 35 cycles of 98°C for 15 s, 52°C for 30 s, 72°C for 18 s and final elongation at 72°C for 5 min. Triplicate PCR reactions (20 μL) using Phusion™ High-Fidelity PCR MasterMix (Thermo Fisher Scientific, USA) were pooled and the amplicons of ∼450 bp were purified from 1.5% agarose (MP Roche, Germany) gel using the Wizzard SV Gel and PCR clean system (Promega, USA) and quantified with Qubit dsDNA HS assay (Thermo Fisher Scientific, USA). Samples were randomly distributed within two runs to account for the batch effect and sequenced on Illumina Miseq 2 x 300 bp PE (Macrogen, South Korea).

Raw reads were quality-checked using FastQC v0.11.7 (Babraham Bioinformatics, Cambridge, UK). The primer sequences were trimmed and read quality filtered using Cutadapt v1.16 maximum error (-e 0.1), quality cut-off (-q 20) and minimum length (-m 250) [58]. Reads were truncated using *filterAndTrim* (truncLen = c(220, 220), maxEE=c(2,5), truncQ=2) in the R/Bioconductor environment from DADA2 package v1.12.1 [59]. ASVs were constructed and chimeric sequences removed using the method “consensus”. ASVs present only in one of the runs were removed from downstream analysis using *intersect* and *subset*. Subsequently, ASVs were aligned in Geneious v2019.2.3 using ClustalW v2.1 [60]. Poorly aligned ASVs were confirmed to not be *puf*M with a blast against NCBI non-redundant database [61] and excluded from further analysis. The final dataset consisted of 1588 ASVs (Supplementary File S3, Reference ASV sheet) and 62,729 ± 13,448 reads per sample (mean ± standard deviation, File S3, ASV_table sheet). The sequences were deposited in the NCBI database under Biosamples SAMN38037304 - SAMN38037518 as a part of BioProject PRJNA970655.

The taxonomic assignment was done through phylogenetic placement using The Evolutionary Placement Algorithm v0.3.5 [62] that placed the ASVs into the phylogenetic tree calculated from the new reference database sequences that were back-translated from protein alignments using trimAl [63]. The taxonomic assignation was handled according to the ASV phylogenetic position using Gappa [64] (Supplementary File S3, Taxonomy sheet).

### Phytoplankton community analysis based on 16S rRNA gene amplicons

The V3-V4 region of the bacterial 16S rRNA gene was amplified using 341F and 785R primer pair [65] as described in Piwosz et al., 2022 [17] The subset of sequences assigned to Chloroplast was extracted and their taxonomy was further affiliated using a curated reference database of the plastidial 16S rRNA gene: PhytoRef [66]. Bar plots were visualized using ggplot v3.4.3 [67].

### Data and statistical analysis

Unless stated otherwise, all analyses were done in R studio v3.6.1 and were visualized using ggplot2 v3.3.6 [67]. Dynamics of environmental and biological variables were interpolated using igraph v1.2.6 and lubridate v1.8.0 [68, 69]. For addressing the compositional bias of amplicon data [70], principal component analysis was done using centred log ratio (CLR) transformation [71] through *transform* from microbiome package v1.17.42. Community composition bar plots and Alphaproteobacteria, Gammaproteobacteria and Gemmatimonadota bubble plots were done using Phyloseq v1.30.0 [72]. The 100 most abundant ASVs were selected and plotted using *plot_heatmap* [72]. The occurrence of specific ASVs in spring was tested using analysis of compositions of microbiomes with bias correction in ANCOMBC v2.3.2 [73, 74] and plotted using ggplot2 v3.4.3 [67].

Relationships between environmental data and AAP community were analysed using distance-based linear models (DistML) [75, 76] in the PERMANOVA+ add-on package of the PRIMER7 software [77] (Primer Ltd., Lutton, UK). From strongly correlated environmental variables (correlation coefficient > 0.6) only one was selected for further analysis. The model was calculated on the CLR-transformed relative abundance data of AAP bacteria [71], using a stepwise selection procedure. The best model was selected based on statistical significance (9999 permutations) and the value of Akaike’s Information Criterion (AICc).

### AAP core and network community analyses

ASVs present in more than 80% of the samples from the 3 years and four depths, were considered the AAP lake core microbiome. The percent contribution of each core ASV to their respective maximum percent contribution was calculated and plotted with bubble plots using Phyloseq v1.30.0 [72] and ggplot2 v3.3.6 [67].

SparCC analysis was applied to calculate lake community co-occurrence correlations from the compositional data [78]. Only correlations with pseudo p-value < 0.02 and stronger correlation than ± 0.7 were selected. The network was plotted using Cytoscape v3.9.1 [79].

### Time series and trend lines

Interannual trend analysis was done in TTR package v0.24.3 [80]. Raw data on total and AAP bacterial abundances, temperature and Chl-*a* concentrations were averaged for months and depths and transformed into time series assuming annual frequency. They were decomposed into trend, seasonal and random components using *decompose* with default settings. Spearman correlation between interannual trends of AAPs abundance and temperature and Chl-*a* concentrations was done for extracted trend component.

## Results

### New database and longer amplicons enhance taxonomic assignments of *puf*M ASVs

In order to improve the taxonomic assignment of the *puf*M gene amplicons, we constructed a new reference database containing 3633 *puf*M sequences (> 646 bp) from Pseudomonadota (synonym for Proteobacteria), Gemmatimonadota, Chloroflexota, Eremiobacteriota and Myxococcota (Supplementary File S1). The database includes 529 genera, 114 families, 53 orders, and 9 classes (Supplementary Figure S1) from cultured species and MAGs originating from a wide variety of habitats (Supplementary File S2), mostly from freshwater (2140 sequences) and marine environments (381 sequences).

The newly designed primer set covers 80.5% of the new database at maximum of three mismatches (Supplementary File S4). The older, highly degenerated primer pairs, UniF + UniR, pufMF + pufMR and UniF + pufM_WAW, cover 98.9%, 85% and 96.2%, respectively. However, the amplicon length of the novel primers is about two times longer (∼450 bp) allowing for proper taxonomic assignation of more than 95% of the alphaproteobacterial and above 75% of the gammaproteobacterial reads at order level. Additionally, 38 alphaproteobacterial, 36 gammaproteobacterial and 6 Gemmatimonadota genera were detected (Supplementary Figures S2, S3 and S4).

### Seasonal changes in Cep lake

Environmental conditions in Cep lake showed seasonal dynamics typical for a temperate freshwater lake (Supplementary File S5). In January and February, the lake was partially frozen and stratified from April/May until September with maximum temperatures >24°C in July and August. The metalimnion was located between 5-8 meter-depth in 2017, and between 2-5-meters in 2018 and 2019. In all three years, the autumnal mixing, characterized by higher values of dissolved oxygen and lower temperatures, was initiated in October (Supplementary Figure S5A-B).

Chlorophyll-*a* measurements varied throughout the year with seasonal maxima representing spring and autumn phytoplankton blooms (Supplementary Figure S5C). The spring phytoplankton bloom terminated at the onset of stratification, and was composed, according to 16S amplicons affiliated to plastids, mostly by Bacillariophyta and Chrysophyceae (Supplementary Figure S6).

### Seasonal dynamics of AAP community composition

The maximal AAP abundances (3.42 - 5.50 × 10^5^ cells mL^-1^) corresponded to 15-20% of the total bacteria (Supplementary Figure S5D-E) and closely proceeded the spring phytoplankton blooms. A second AAP bacterial peak occurred towards the end of summer, before the autumn phytoplankton peak. Alpha diversity of AAP community was lower during the spring abundance peaks and rose during the second part of the year (Supplementary Figure S5F). The AAP community followed an annual recurrent pattern during the three consecutive years, with a distinction between the epilimnion and hypolimnion communities during the stratified period (Figure 1A). Samples from the same season in different years were more similar to each other than samples from different seasons in the same year, indicating the persistent temporal succession of the community and similar interannual community structure. Distance-based linear models (DistLM) and distance-based redundancy analysis (dbRDA) selected temperature, Chl-*a*, and total, Cyanobacterial, and AAP abundances, to best explain the variability (23.11%) of the AAP community composition (Supplementary File S6). For 2018 and 2019, two years for which nutrient data is available (Supplementary File S5), phosphorous and ammonia increased this explanation by 6.75%, up to 29.86%. Interestingly, in addition to a seasonal cycle, we observed an interannual variation in AAP community composition (Figure 1B). The decomposition of time series on monthly averaged values for the whole water column showed increasing interannual trends in temperature and AAP abundance during 3 years of sampling, and a decreasing trend in Chl-*a* concentration (Supplementary Figure S7). Trends of temperature and AAPs abundance were significantly correlated (Spearman correlation coefficient rho = 0.8, p-value < 0.0001).

**Figure 1:**
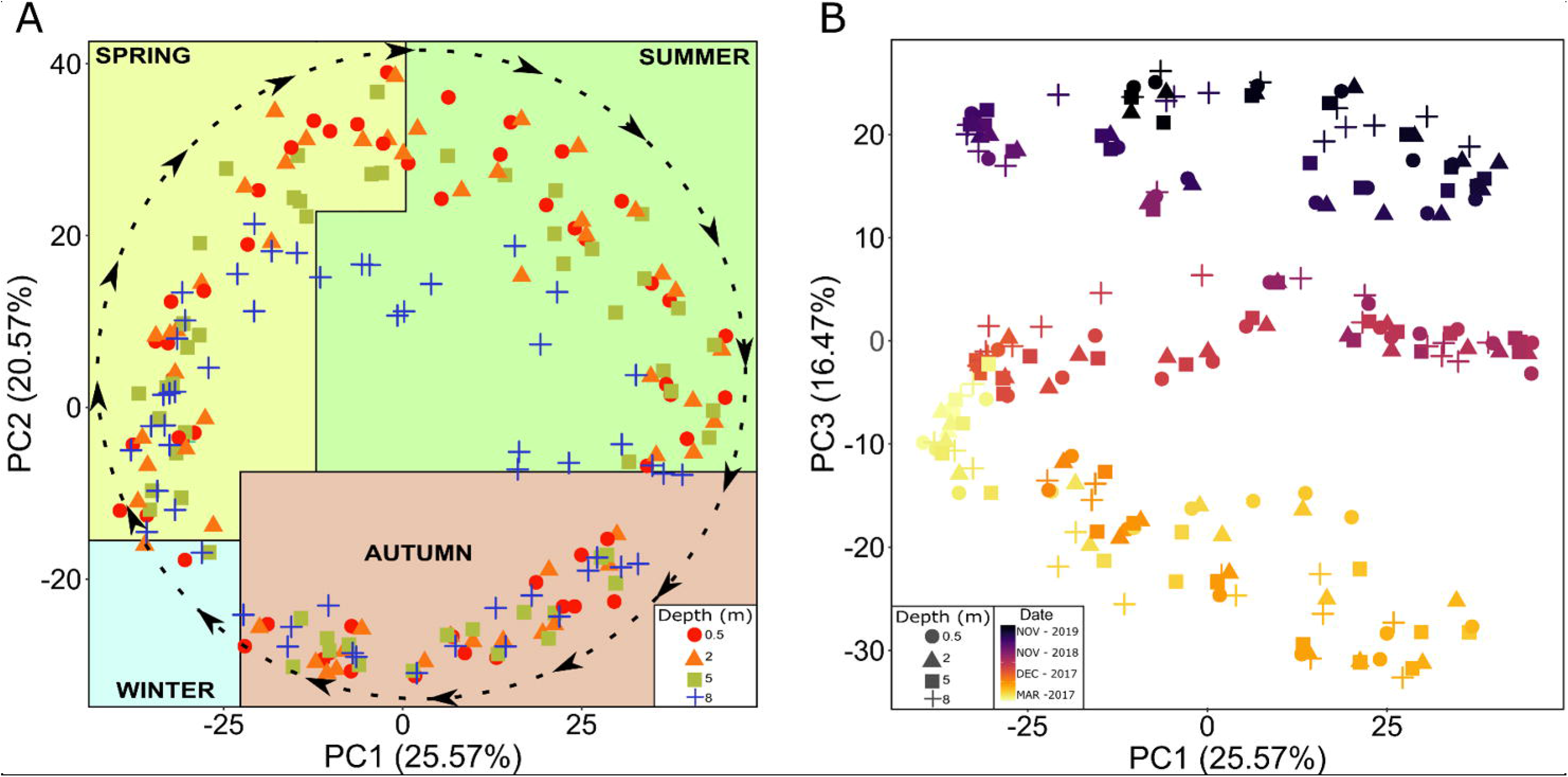
Development of AAP community structure. Principal component analysis of centred log-ratio transformed AAP community composition. Each point represents a sample with 0.5 m (red circle), 2 m (orange triangle), 5 m (green square) and 8 m (blue cross) in PC1 and PC2 axis (A) and 0.5 m (circle), 2 m (triangle), 5 m (square) and 8 m (cross) coloured according to the date of sampling in PC1 and PC3 axis (B). Dashed line and arrows in panel A indicate AAP community succession following an annual chronological direction.

The AAP community was dominated by Gammaproteobacteria over the whole water column, with an average relative contribution exceeding 50% and reaching up to 90% during stratification (Supplementary Figure S8). Alphaproteobacteria was the second most abundant class, showing the maxima contributions in spring and autumn, reaching over 50% in the spring of 2018. Classes Gemmatimonadetes (Gemmatimonadota) and Myxococcia (Myxococcota) made up 5% and 2% of the AAP community, respectively. Unclassified Pseudomonadota and unclassified Myxococcota showed transient contributions of <9% and <1% of the AAP community, respectively. Chloroflexota and Eremiobacteriota were not detected.

The 100 most abundant ASVs (based on their average relative abundances) comprised 75 % of the reads and exhibited seasonal recurrence, peaking every year at specific times of the year (Figure 2). The majority of ASVs demonstrated a transient contribution and were generally absent outside their maxima (e. g. ASV28 (Rhizobiales)), while a few showed pronounced relative abundance throughout the year (e. g. ASV4 (*Rhodoferax*)). Interestingly, ASV67 (*Limnohabitans*), whose relative abundance was on average <0.36%, was the sole ASV detected in every sample.

**Figure 2:**
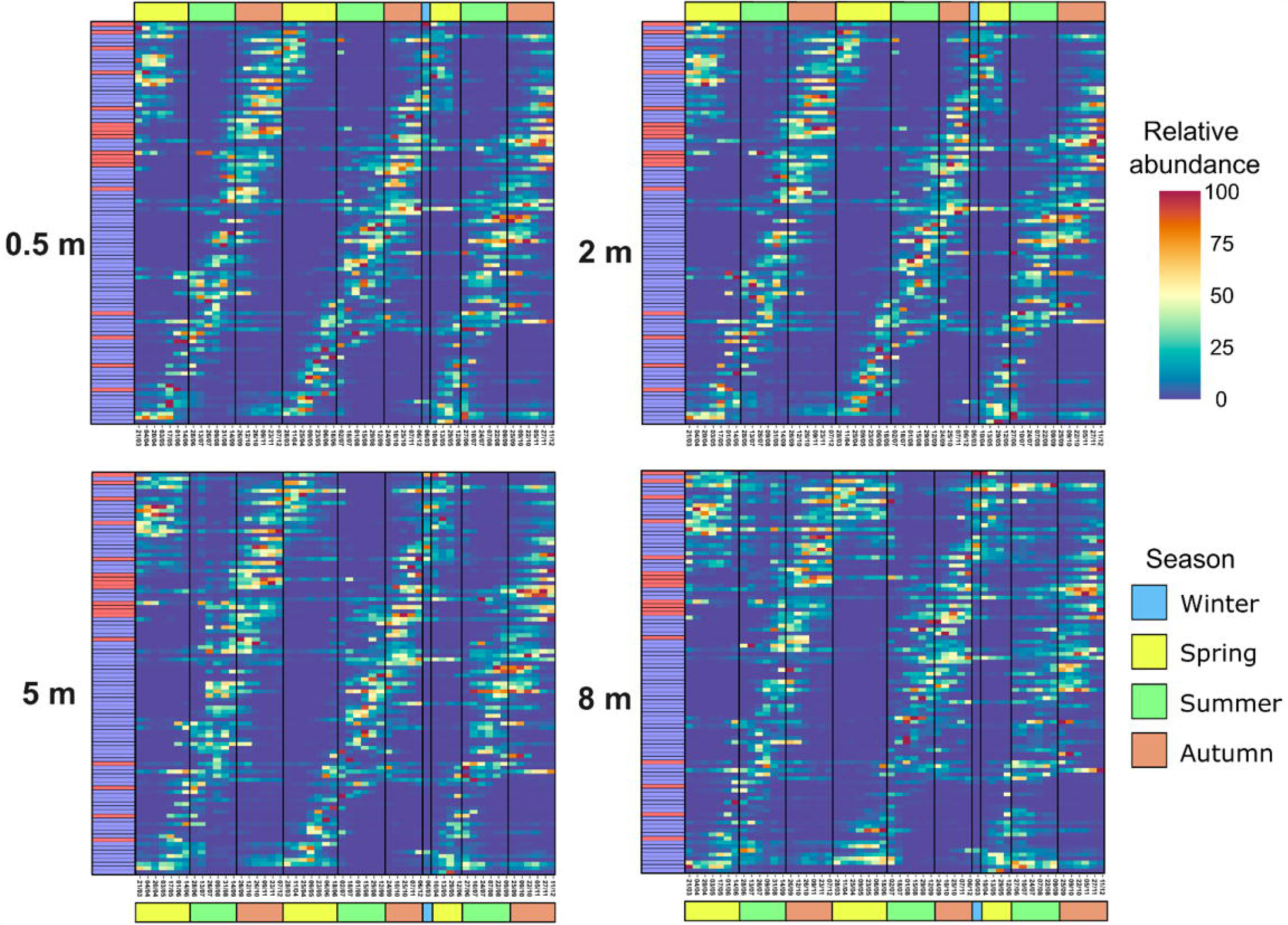
Recurrence of 100 most abundant ASVs. Relative abundance is individually normalised in 0.5 m, 2 m, 5 m and 8 m during the different seasons of the 3-year sampling. Coloured box on the left indicate the taxonomic assignation at class level of each ASV, red for Alphaproteobacteria and blue for Gammaproteobacteria. Colour bar on the top and bottom indicate the seasons of the year.

Out of 1588 *puf*M ASVs (Supplementary File S3, Reference ASV sheet), the stable part of the AAP community (defined here as ASVs present in >80% of the samples) consisted of only 22 ASVs (Supplementary Figure S9): 8 *Rhodoferax*, 4 *Limnohabitans*, 2 *Aestuariivirga*, 1 *Methylobacterium*, 1 *Rubrivivax* and 6 other Burkholderiales. These core ASVs varied largely in their contribution from the most abundant ASV2 (*Aestuariivirga*, with an average relative abundance of 4%) to the least abundant ASV237 (unclassified Burkholderiales, with an average relative abundance of 0.06%). Their seasonal dynamics differed substantially throughout the year and distinct relative abundance patterns were observed even for ASVs from the same genus. For instance, ASV5 and ASV49, both *Rhodoferax*, peaked in autumn and spring, respectively. Similar differences were observed for two *Aestuariivirga*: ASV2 peaked during the spring mixing period (from March to May), while ASV62 showed its highest contribution during the summer stratification. It is noticeable that the core AAP community also included ASVs outside the 100 most abundant, such as ASV115 (*Rhodoferax*) and ASV237 (unclassified Burkholderiales), which had a low but steady contribution during the whole sampling season. Furthermore, dynamics of some core ASVs, such as ASV31, were different each year.

To identify phylotypes most contributing to the difference in AAP community composition during spring and autumn peaks, we carried out an analysis of compositions of microbiomes with bias correction (Supplementary File S7). Composition of genera and orders contributing to the AAP bacterial peaks was different (Figure 3). Spring peak consisted of a higher prevalence of Alphaproteobacteria versus Gammaproteobacteria genera (9 vs 5), while during autumn, the community was more diverse and included also Gemmatimonadota and Myxococcota. The highest genera contributors to the spring peak (log fold change > 2) were *RFPW01*, *UBA1936*, *Cypionkella* and *Rhodoferax*, while *UBA964* and *UBA5518* (Gammaproteobacteria: Steroidobacterales and Pseudomonadales, respectively) contributed most to the late summer peak. The only genera substantially contributing in both peaks was *Aestuariivirga*.

**Figure 3:**
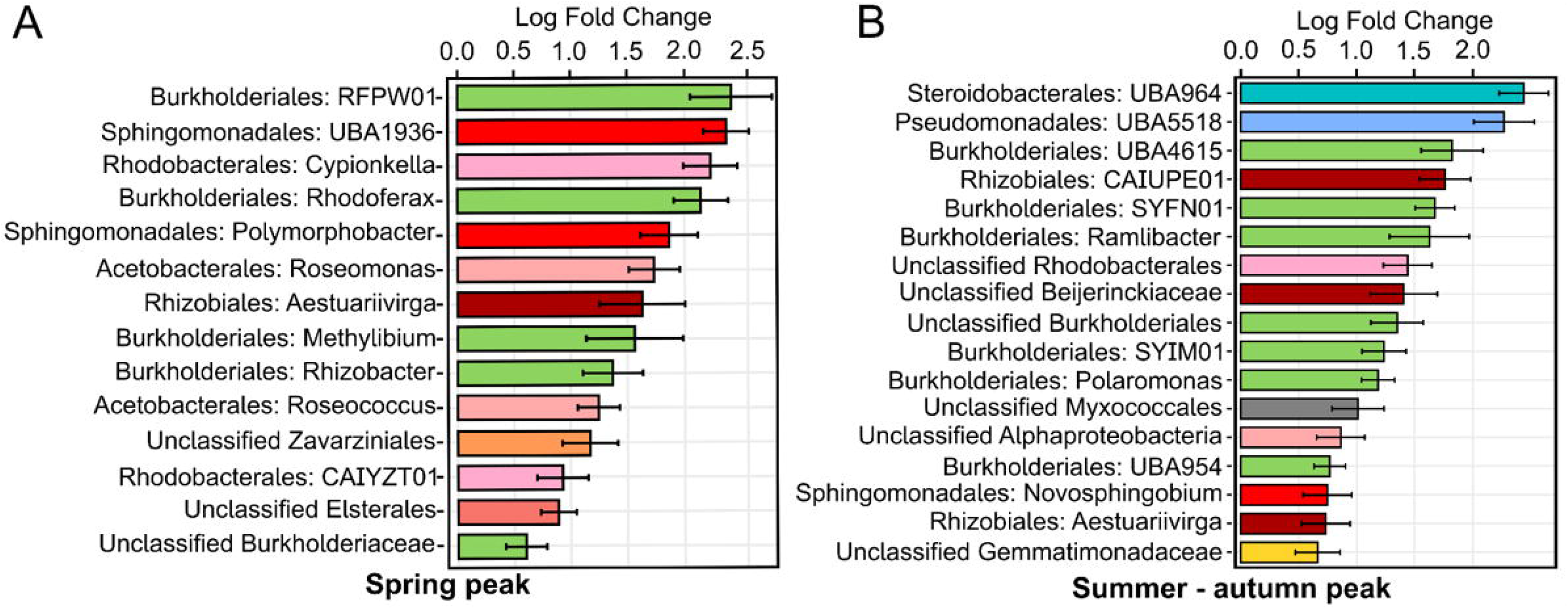
Community composition of AAP abundance peaks. Analysis of compositions of microbiomes showing log fold change values at order and genus level in Spring (A) and in Summer-Autumn peaks (B) with green colour scale for Burkholderiales, blue colour scale for other Gammaproteobacteria, red colour scale for Alphaproteobacteria, yellow for Gemmatimonadota and grey for Myxococcota orders.

### Network analysis

To study possible interactions between AAP bacteria, we performed a network co-occurrence analysis. The network calculated for the entire lake included 99 ASVs and 139 interactions (92% positive; Figure 4). Thirteen highly connected nodes with more than six co-occurrence correlations were identified as hubs. They belonged to genus UBA964, Steroidobacterales, *Rubrivivax* and unclassified Burkholderiaceae from Gammaproteobacteria. The most connected nodes from Alphaproteobacteria belonged to *Aestuariivirga* and Rhodobacteraceae (4 edges each). The majority of correlations (∼60%) occurred between ASVs from the same genus, family or order, creating groups of densely connected nodes (e.g. genus UBA964). Furthermore, some ASVs peaking in spring correlated positively with each other (*Aestuariviirga* and unclassified Burkholderiaceae) in contrast to *Rhodoferax* and *Cypionkella*. A similar pattern was observed for ASVs peaking in summer-autumn.

**Figure 4:**
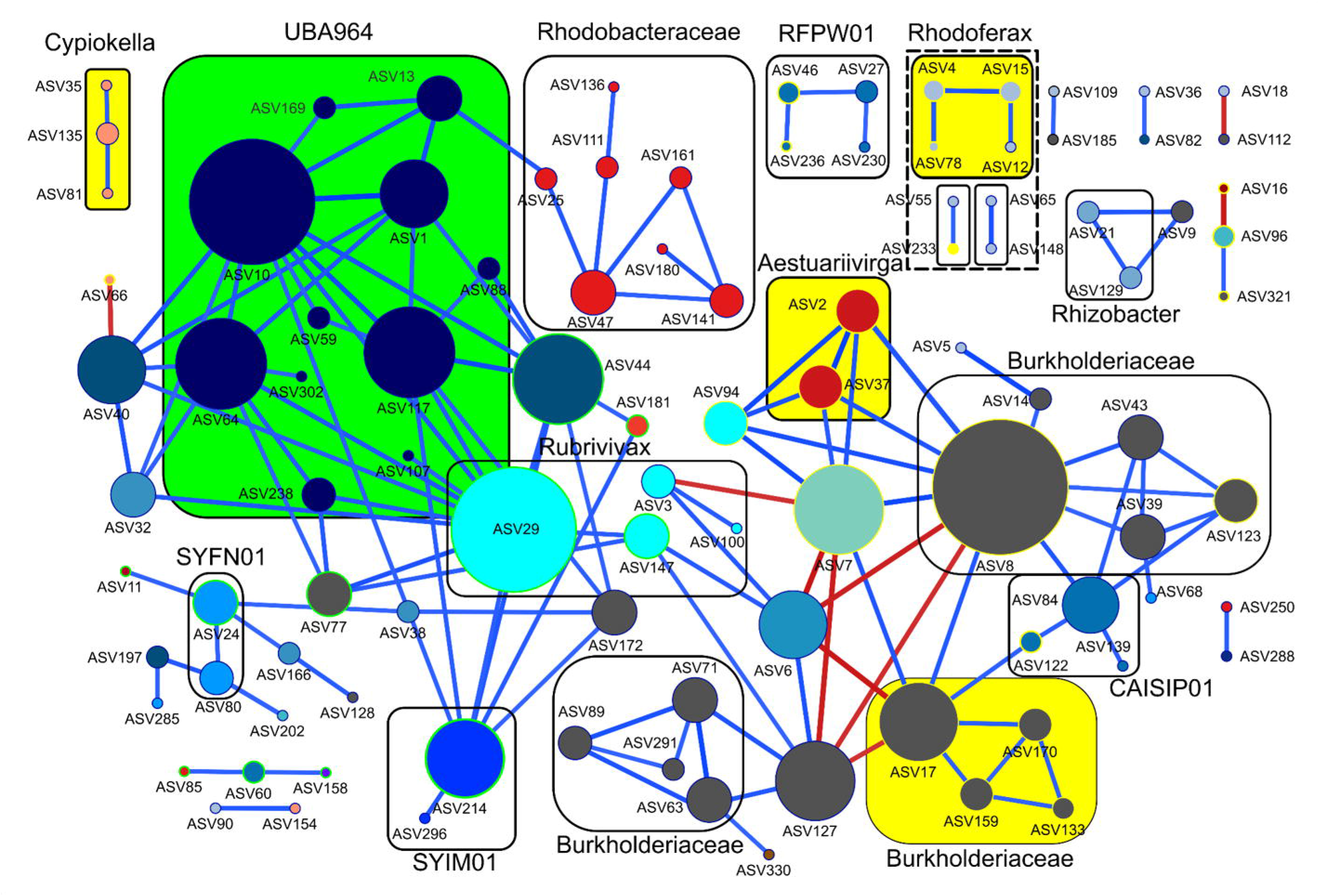
AAP community co-occurrence correlation network. The radius of the nodes (ASVs) are directly proportional to the number of significant correlations. Colour of the nodes shows the taxonomy at class level with Alphaproteobacteria in red-orange colour scale, Gammaproteobacteria blue colour scale, unclassified Burkholderiaceae (grey) and Gemmatimonadota (brown). Blue lines indicate positive and red lines negative correlations. Rectangles with solid lines indicate nodes that interact with other nodes from the same taxonomic rank (genus or family) and positive log fold change for Spring AAP peak are highlighted in yellow rectangles fill and yellow node stroke while green rectangles fill and green node stroke for Autumn AAP peak.

## Discussion

Seasonal succession of planktonic communities in temperate lakes has been intensively studied [2]. The PEG model defines the key phases in the annual development of ecological succession as well as the interactions between different organisms in aquatic ecosystems. The annually recurrent phenomena in freshwater lakes include a spring phytoplankton bloom followed by a zooplankton-induced clear-water phase in early summer, a late-summer phytoplankton bloom and a period of low productivity in winter. Recently, seasonal dynamics of heterotrophic bacteria have been incorporated into the model [5], as they rapidly respond to the transitions in the lakes’ pelagic functioning, especially during the phytoplankton bloom [3, 5, 81]. Whilst AAP bacteria have been shown to substantially contribute to the bacterial abundance, biomass and activity in freshwater lakes [13, 17], they are not considered in the PEG model [1, 2].

Dynamics of microbial communities are typically investigated by sequencing of the 16S rRNA gene and such analyses do not allow for disentangling the metabolic functionality of bacteria, especially when traits of interest follow a heterogenic pattern of presence within the same taxonomic ranks. This is the case for AAP bacteria, where members from the same genus might or might not contain the ability to carry out anoxygenic photosynthesis [26]. Amplicon analysis of a functional gene may overcome this hindrance, but it requires a comprehensive database with taxonomically assigned reference sequences. Currently, some phylogenetic clades (A-L) cannot be assigned to the genus or even order level [82, 83]. Moreover, different researchers assemble databases for their environment of interest, with different quality thresholds and criteria [17, 84], which hampers direct comparison between different studies. Finally, short amplicons obtained with the most commonly used primer combinations, puf_UniF - puf_UniR, puf_UniF – pufM_WAW and pufMF – pufM_WAW [27, 32] often do not allow for taxonomic assignment below the class or order level, resulting in a substantial number of unclassified reads [18, 29, 84, 85].

### Comprehensive database and longer amplicons allow for improved taxonomic assignments of *puf*M ASVs

We constructed the largest curated *puf*M database to date that includes sequences from essentially all environments, with especial high representation of freshwater lakes (Supplementary Figure S1, Supplementary File S1 and S2). The key improvement is that we included only sequences originating from taxonomically assigned genomes and MAGs, excluding all environmental *puf*M sequences classified into phylogroups based on phylogenetic analysis [82]. The reason for this exclusion was that phylogenetic trees based on *puf*M gene do not match 16S rRNA phylogeny due to horizontal gene transfer events [86–88]. In contrast, taxonomic assignment of genomes and MAGs based on whole genome or 120 selected marker genes is more accurate and consistent [89]. During our quality control, sequences originating from Bdellovibrionota, Verrucomicrobiota, Omnitrophota, Planctomycetota or Bacteroidota were excluded from the final database. Some members of these phyla were reported to encode *puf*M [85]. However, our manual inspection revealed that their *puf*M gene was present in short contigs, and often found as the only phototrophic gene, which does not warrant phototrophic functionality. Furthermore, *puf*M genes from these phyla did not form a monophyletic clade, suggesting dubious multiple and independent events of horizontal gene transfer. Thus, only members of Pseudomonadota, Chloroflexota, Gemmatimonadota, Eremiobacteriota and Myxococcota were included. Our choice of sequences ensures high quality and allows for future extensions of the database as more metagenomic data is produced and more AAP bacteria are cultured, enhancing its fidelity and functionality. Moreover, as our database contains entire or almost entire *puf*M gene sequences, it can be used for amplicon taxonomy assignment independently of the actual primers used.

The *puf*M gene is one of the most conserved genes from the *puf* operon which codifies the genes for synthesis of the anoxygenic photosynthesis apparatus [90, 91]. Commonly used puf_UniF - puf_UniR, puf_UniF – pufM_WAW and pufMF – pufM_WAW primer pairs [27, 32] hybridize on the most conserved regions at the end of the gene, separated by ∼110 - 160 bp. They show high coverages (Supplementary File S4) but produce short amplicons that hamper taxonomic assignation below the class or order level resulting in a high fraction of unclassified reads [18, 29, 84, 85]. Thus, we designed a new primer set producing longer amplicons to increase the taxonomic resolution, as has been shown for other genes [92]. The new primer pair has lower *in silico* coverage against our new database than puf_UniF - puf_UniR [27] (80.5 vs 98.9 %). Some groups, such as Chloroflexota, *Aquidulcibacter* and *Polynucleobacter*, were poorly covered, which may explain their absence in our amplicons. Nevertheless, the number of sequences identified at every taxonomic level increased compared to a previous study in the same lake [18]: 95% of the alphaproteobacterial and above 75% of gammaproteobacterial reads were classified at the order level. Additionally, the number of newly detected genera was substantially higher (80 vs 12) and the Shannon index showed a wider range of diversity (Supplementary Figure S5F), since longer amplicons enable to detect more nucleotide variations, and thus reveal higher diversity, advancing our knowledge on AAP community composition.

### Phenology of AAP bacteria and consideration into the PEG model

The seasonal succession of the AAP community revealed differential strategies of adaptation to environmental conditions, unveiling generalist AAP bacteria appearing most of the time, whereas specialists or opportunists showed a transient contribution to the AAP community (Figure 2). Within the generalists, we identified the core AAP community that consistently contributed throughout the seasons for the three consecutive years and across all depths (Supplementary Figure S9). The coexistence of the core AAP community, the dominance of positive correlations in the networks and the numerous correlations between ASVs of similar taxonomic ranks (Figure 4), suggest partial metabolic redundancy within some closely related AAP bacteria that maintain functional kinship. In contrast, the complex seasonal succession pattern indicates that the lake’s AAP community is extremely diverse. AAP community represent a large functional repertoire (even within the same genus) allowing for niche speciation *via* temporal succession, facilitating their geographical coexistence. Finally, the 3-year recurrence of the AAP community (Figure 1A) documents its indigenous character in freshwaters and the higher importance of selection over the environmental drift and dispersal processes at short temporal scale [93]. Changes in the AAP abundance coincided with shifts in their community composition indicating that abundance peaks were caused by specific phylotypes. These phylotypes differed between both abundance peaks (Figure 3). Generally, AAP community was dominated by Gammaproteobacteria except for spring abundance peaks, when Alphaproteobacteria, that already have shown higher phototrophic activities in spring [31], increased their contribution. Additionally, the directional interannual variation of the AAP community (Figure 1B) signifies the evolution of AAPs populations, potentially influenced by changes in environmental and biological variables such as temperature or Chl-*a* (Supplementary Figure S7).

While the PEG model has enhanced our understanding of seasonal patterns, it still does not encompass all aquatic components, such as viruses or specific functional bacterial groups. This includes AAP bacteria, which are characterized by a heterogenic behaviour but still represent an important functional group, fulfilling valuable ecological and biochemical processes in the aquatic environment. For that reason, amending them into the present PEG model will certainly improve our understanding of aquatic community functioning.

Cep lake is representative of a common temperate lake in the northern hemisphere: it is small, shallow and meso-oligotrophic [94], thus our conclusions might be applied to other lakes. AAP bacteria play an important role during and shortly after the spring bloom when their abundance and contribution to the total bacterial community were recurrently the highest. This spring AAP abundance peak preceded those from the overall heterotrophic bacteria (Figure 5). The faster response of AAP bacteria highlights that photoheterotrophy confers a distinct metabolic advantage that triggers their growth in this scenario over other heterotrophic bacteria. Their importance as food for bacterivores in microbial food webs is well documented [95–97] and due to their, on average, larger cell size and higher activity than other heterotrophic bacteria [13, 14], they contribute disproportionally to the carbon cycling contrary to their relatively low abundances [19, 97]. Additionally, Chl-*a* concentration has been identified as a variable explaining the dynamics of the AAP bacterial community and it is plausible to assume that the spring abundance peaks of AAP bacteria are triggered by the excess of carbon released by the phytoplankton bloom (mostly diatoms; Supplementary Figure S6) and the lack of grazing pressure after winter. Moreover, AAPs have a highly efficient photoheterotrophic metabolism [17] increasing secondary bacterial production and disposing more carbon to higher trophic levels *via* the microbial loop. This emphasizes the urgent need for more quantitative studies to further decipher carbon transfers along microbial and classical food webs. The AAP bacterial peak is terminated by selective and extensive grazing of bacterivorous protists and macrozooplankton that are also present in summer [81, 96, 98]. The absence of a pronounced AAPs peak following the phytoplankton bloom in autumn (Figure 5) could be attributed to the higher grazing pressure, distinct phytoplankton composition (Supplementary Figure S6), decreasing temperatures (Supplementary File S6), and the decreasing light availability at shorted day length [2, 18, 30, 99].

**Figure 5:**
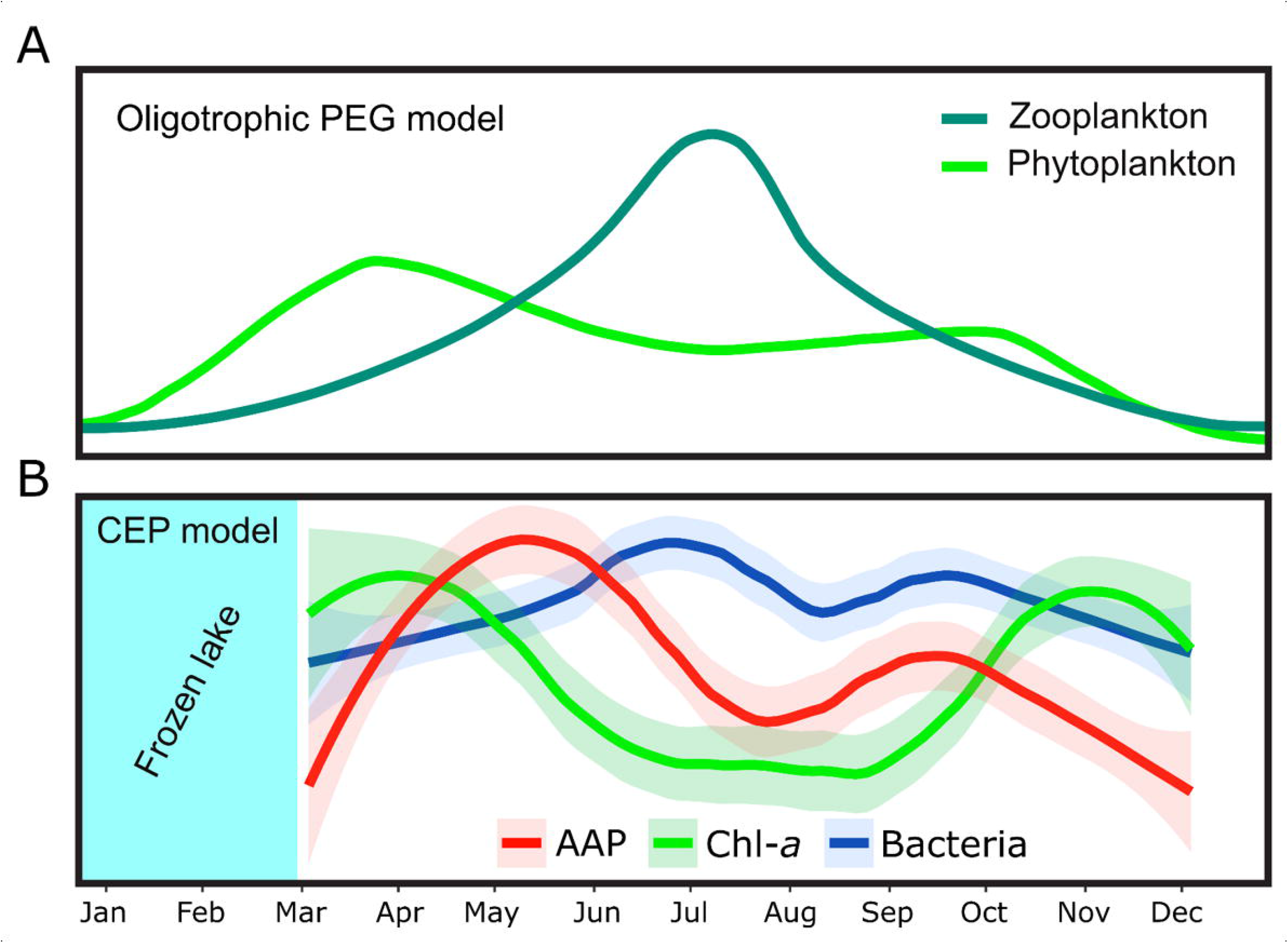
Including AAP bacteria in the PEG model. Annual succession patterns of microbial communities for (A) Phytoplankton and zooplankton according to original PEG model in oligotrophic scenario, and (B) monthly averaged annual succession pattern of AAP abundance, trophic status through Chl-a, and of bacterial abundance. Trend lines are normalized to maxima and minima values for each variable.

Finally, our study provides novel insight into the ecology of phototrophic Myxococcota. While their average contribution was low, they were recovered during stratification over three consecutive years (Supplementary Figure S8) and constituted a member of the summer-autumn AAPs peak, emphasizing their potential significance in microbial communities during summer as they showed a potentially predatory and photoheterotrophic metabolism[100].

## Conclusions

Our study revealed annual recurrent seasonal patterns of AAP bacteria in a freshwater lake, supporting the conceptual inclusion of this important functional group into the PEG model. The high abundance of AAP bacteria during the spring phytoplankton bloom highlights their crucial role in recycling phytoplankton-derived dissolved organic matter and their role in aquatic food webs, which needs to be further quantified and better understood. Differential contribution patterns of the core community and temporal succession of the AAP community indicate strong competition within AAP bacteria communities, which forces them to conduct temporal niche partitioning in order to geographically coexist. In contrast, positive co-occurrence correlations between closely related AAP bacteria indicated their functional redundancy. Our findings provide unprecedented insights into the phenology of AAP bacteria in temperate freshwater ecosystems.

## Conflict of interest

All authors declare no conflict of interest.

## Author’s contributions

CVA, KP and MK did the conceptualization; CVA, IM, KP, JW, AA, CRG, MS, HJR, SS, VK, SA, HPG and RG curated or provided data for *puf*M database; formal analysis was carried out by CVA, KP, LKF and RG; investigation and experimental processes were done by CVA, IM and KP; methodology of database was developed by RG and CVA; CVA and KP wrote original draft and all the authors helped during the review and editing of the manuscript; JD and MH performed the sampling and measured of environmental variables.

## Acknowledgements

The authors want to thank Jürgen Tomasch, Esther Rubio Portillo and Iva Stojan for the insights into the code for data analysis. The Limnos metagenomic dataset was enabled by technical and financial support of the technician team at IGB Stechlin and the Leibniz foundation, respectively.

## Supplementary Figures

<u>Supplementary Figure S1:</u> Maximum likelihood phylogenetic tree of pufM gene sequences of the constructed database. Outer ring represents the environment of origin and the colour of the clades between branches and the outer ring shows the taxonomic classification of the sequences at class level. Colour of the branches refers to the ultra-fast bootstrap values.

<u>Supplementary Figure S2:</u> Alphaproteobacteria community composition at order and genus level for 3-year sampling at 0,5 (A), 2 (B), 5 (C) and 8 m depth (D). Larger size and brighter colours are directly proportional to the relative contribution of each genus to the total Alphaproteobacteria community.

<u>Supplementary Figure S3:</u> Gammaproteobacteria community composition at order and genus level for 3-year sampling at 0,5 (A), 2 (B), 5 (C) and 8 m depth (D). Larger size and brighter colours are directly proportional to the relative contribution of each genus to the total Gammaproteobacteria community.

<u>Supplementary Figure S4:</u> Gemmatimonadota community composition at order and genus level for 3-year sampling at 0,5 (A), 2 (B), 5 (C) and 8 m depth (D). Larger size and brighter colours are directly proportional to the relative contribution of each genus to the total Gemmatimonadota community.

<u>Supplementary Figure S5:</u> Environmental and biological variables for 8 meters’ depth profile during 3-year sampling in CEP lake. Temperature (A), AAP abundance (B), dissolved oxygen (C), percentage contribution to total bacterial community (D), Shannon alpha diversity values (E), and Chlorophyll-a (F). Light-blue vertical bands represent lack of sampling due to frozen lake surface.

<u>Supplementary Figure S6:</u> Phytoplankton chloroplast-based community composition at class level for 0,5, 2, 5 and 8 meters’ depth during 3-years temporal series.

<u>Supplementary Figure S7:</u> Decomposition of additive time series for bacterial abundance (a), AAP abundance (b), chlorophyll-a concentration (c) and temperature (d). Analysis was done in the TTR package version 0.24.3 (R version 4.2.0). Spearman correlation of the decomposed trends between AAP abundance and chlorophyll-a (e) and AAP abundance and temperature (f). R: spearman’s rho value, p: p-value.

<u>Supplementary Figure S8:</u> AAP bacteria community composition according to pufM gene taxonomic assignment at class level for 0,5 (A), 2 (B), 5 (C) and 8 meters’ depth (D) during 3-years sampling campaign

<u>Supplementary Figure S9:</u> Individually normalized relative abundance of the 22 core AAP ASVs during 3 years in 4 depths. Brighter colours and bigger dots indicate larger contribution to the AAP bacterial community. ASVs are clustered according to taxonomic classification at the maximum possible level (genus, family or order).

## Supplementary Files

<u>Supplementary File S1</u>: Nucleotide pufM gene sequence of the database in fasta format.

<u>Supplementary File S2</u>: ID of the pufM gene sequences from the database and their environment of origin.

<u>Supplementary File S3</u>: File containing all the information from the amplicon sequence analysis. Reference ASVs (IDs and sequences), ASV table (ID and abundance on each sample) and Taxa (ID and taxonomic assignation of each ASV).

<u>Supplementary File S4</u>: Primer coverage comparison of the most commonly used pufM gene primer pairs and the newly designed one with 0, 1, 2 and 3 mismatches (MM). Numbers represent the percentage of sequences from the pufM gene database covered for different phyla and classes.

<u>Supplementary File S5</u>: Sample identification number (Sample name) and all the environmental and biological variables measured.

<u>Supplementary File S6</u>: Draftsman plot correlation of the environmental and biological variables, samples removed due to lack of environmental variables, marginal and sequential test for DistLM for 3 years and 2 years (includes also nutrients).

<u>Supplementary File S7</u>: Positive log fold change (lfc) values, standard error (SE) at genus and ASV levels for the spring and autumn AAP abundance peaks with the p- and q-values (p, q).

## Notes

### Competing Interest Statement

The authors have declared no competing interest.

