## Supplementary Figure S1. for "Phenology and ecological role of Aerobic Anoxygenic Phototrophs in fresh waters"

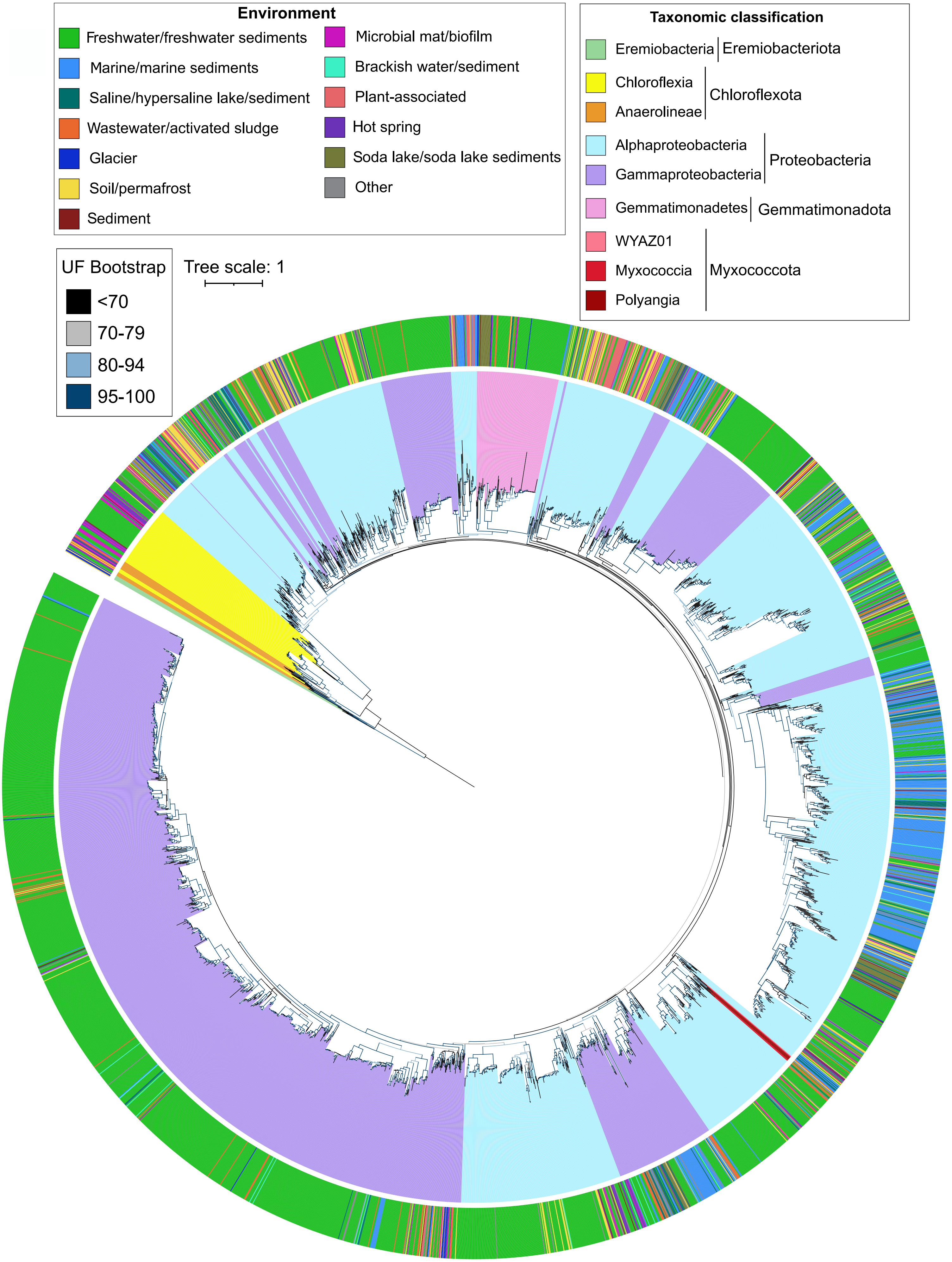


Supplementary Figure S1: Maximum likelihood phylogenetic tree of *puf*M gene sequences of the constructed database. Outer ring represents the environment of origin and the colour of the clades between branches and the outer ring shows the taxonomic classification of the sequences at class level. Color of the branches refers to the ultrafast bootstrap values.
