## Supplementary Figure S3 for "Phenology and ecological role of Aerobic Anoxygenic Phototrophs in fresh waters"

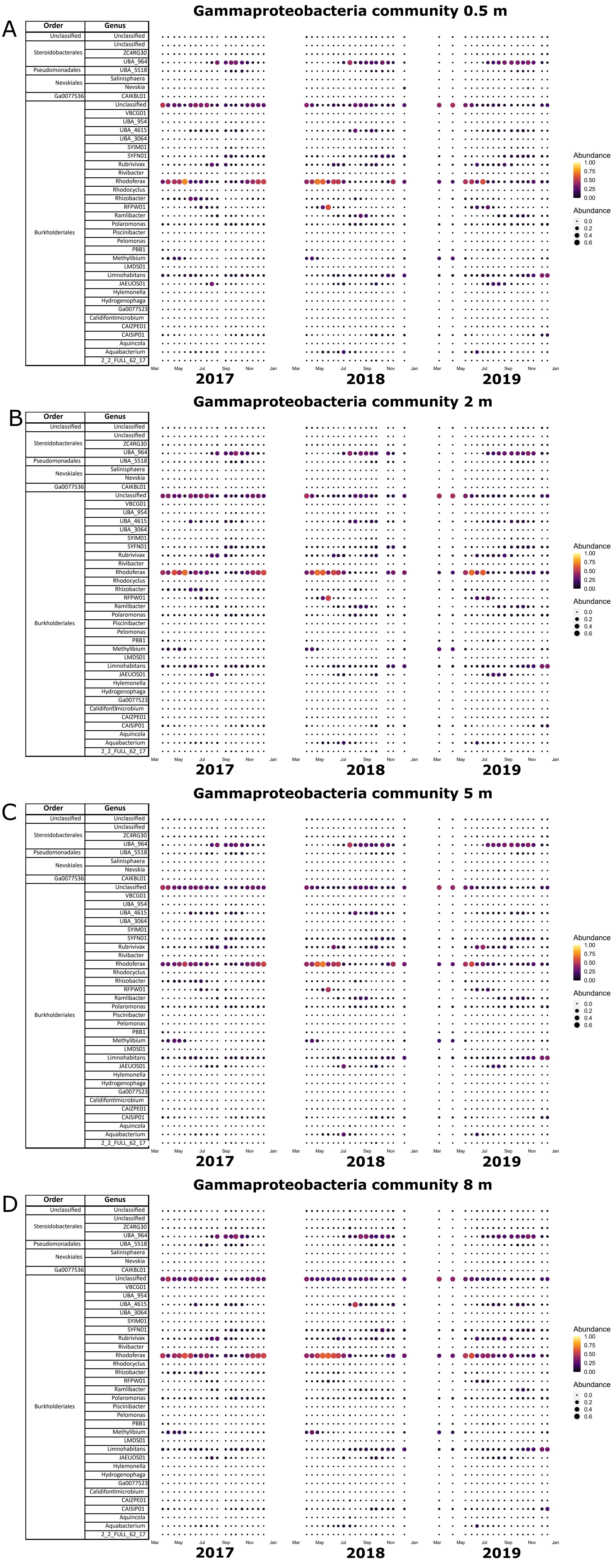


Supplementary Figure S3: Gammaproteobacteria community composition at order and genus level for 3-year sampling at 0.5 (A), 2 (B), 5 (C) and 8 m depth (D). Larger size and brighter colours are directly proportional to the relative contribution of each genus to the total Gammaproteobacteria community
