## Supplementary Figure S5 for "Phenology and ecological role of Aerobic Anoxygenic Phototrophs in fresh waters"

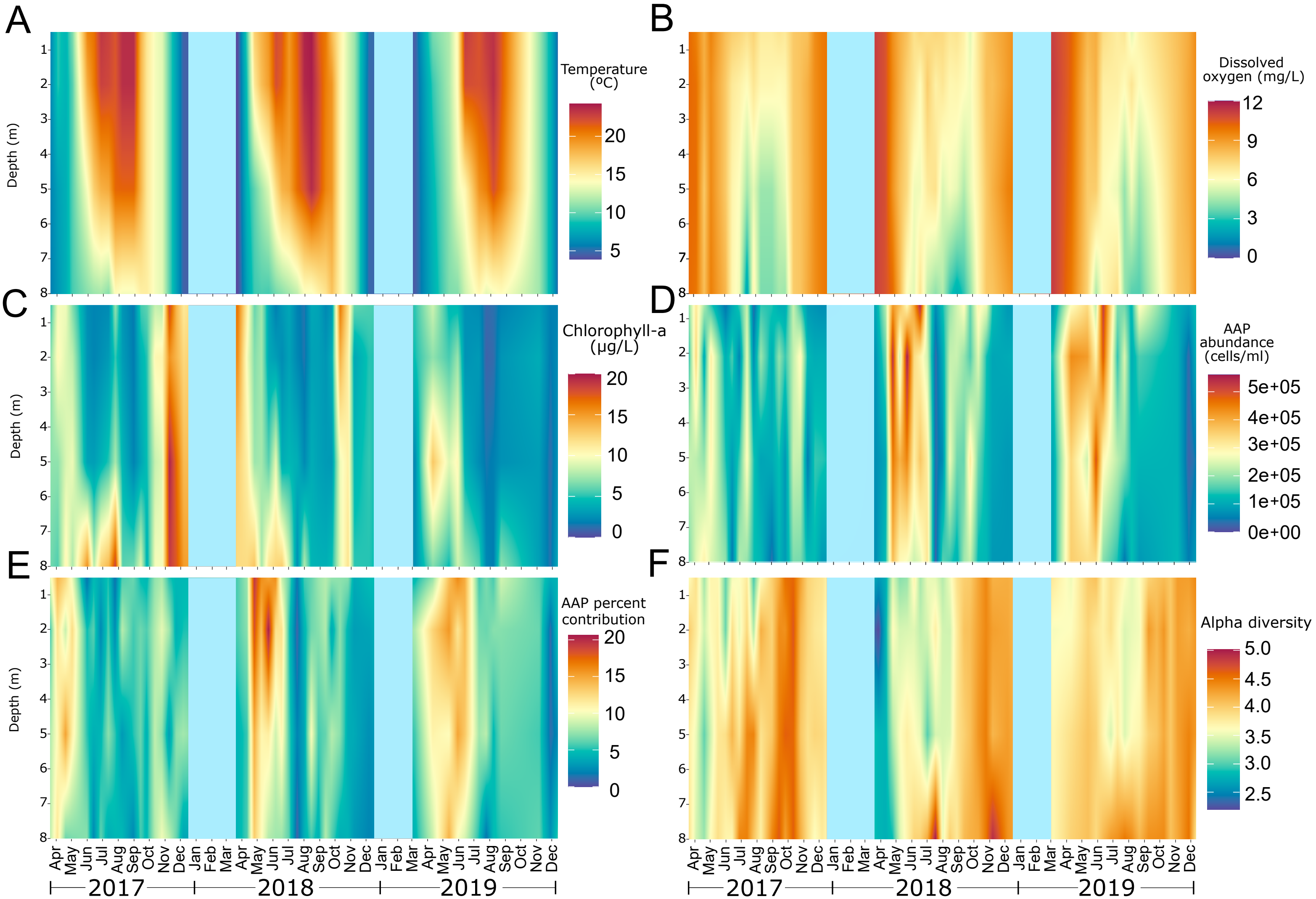


<u>Supplementary Figure S5:</u> Environmental and biological variables for 8 meters' depth profile during 3-year sampling in CEP lake. Temperature (A), AAP abundance (B), dissolved oxygen (C), percentage contribution to total bacterial community (D), Shannon alpha diversity values (E), and Chlorophyll-*a* (F). Light-blue vertical bands represent lack of sampling due to frozen lake surface.
