## Supplementary Figure S6 for "Phenology and ecological role of Aerobic Anoxygenic Phototrophs in fresh waters"

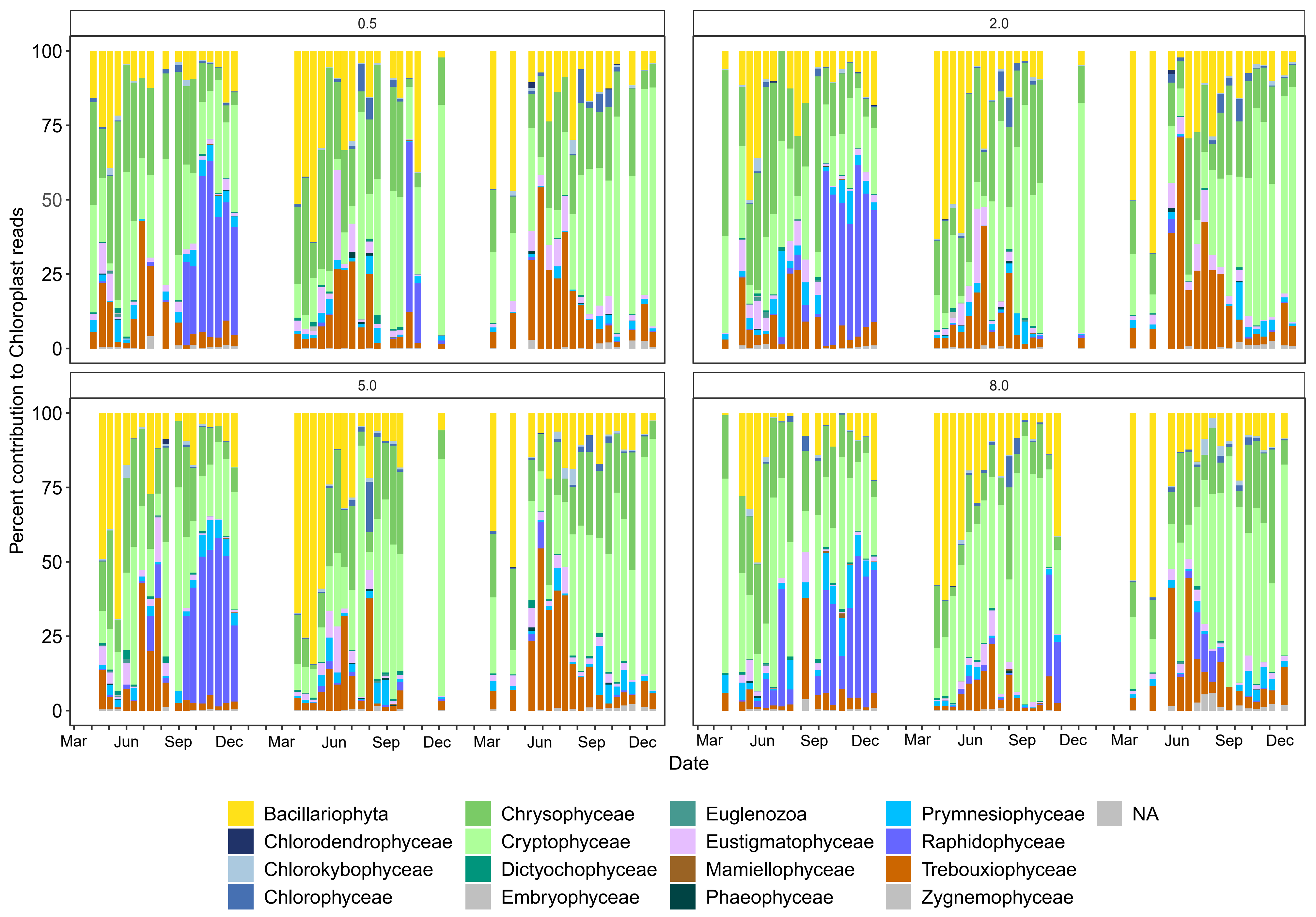


Supplementary Figure S6: Phytoplankton chloroplast-based community composition at class level for 0.5, 2, 5 and 8 meters' depth during 3-years temporal series.
