## Supplementary Figure S7 for "Phenology and ecological role of Aerobic Anoxygenic Phototrophs in fresh waters"

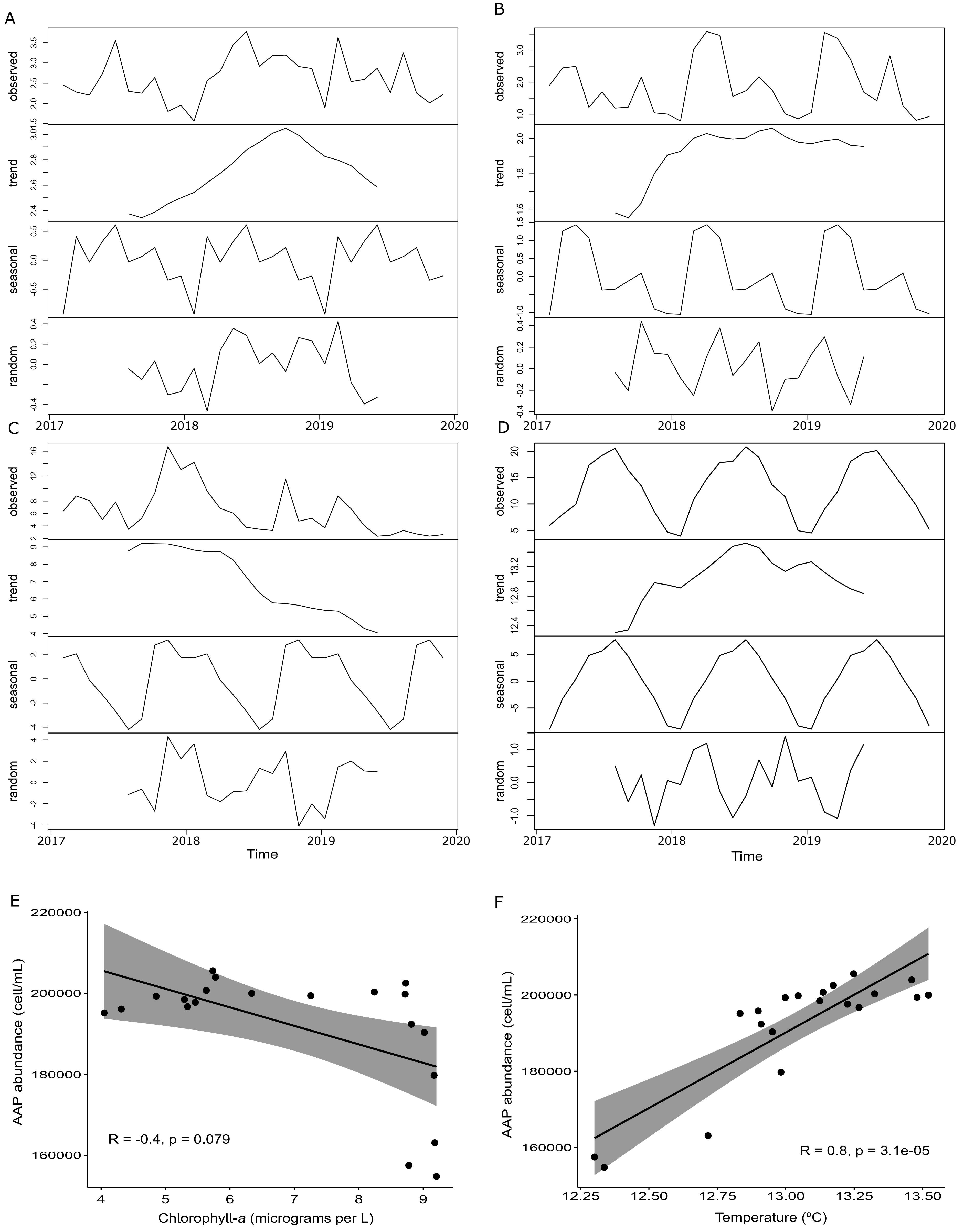


Supplementary Figure S7 : Decomposition of additive time series for bacterial abundance (A), AAP abundance (B), chlorophyll -*a* concentration (C) and temperature (D). Analysis was done in the TTR package version 0.24.3 (R version 4.2.0). Spearman correlation of the decomposed trends between AAP abundance and chlorophyll-a (E) and AAP abundance and temperature (F). R: spearman's rho value, p: p-value.
