## Supplementary Figure S8 for "Phenology and ecological role of Aerobic Anoxygenic Phototrophs in fresh waters"

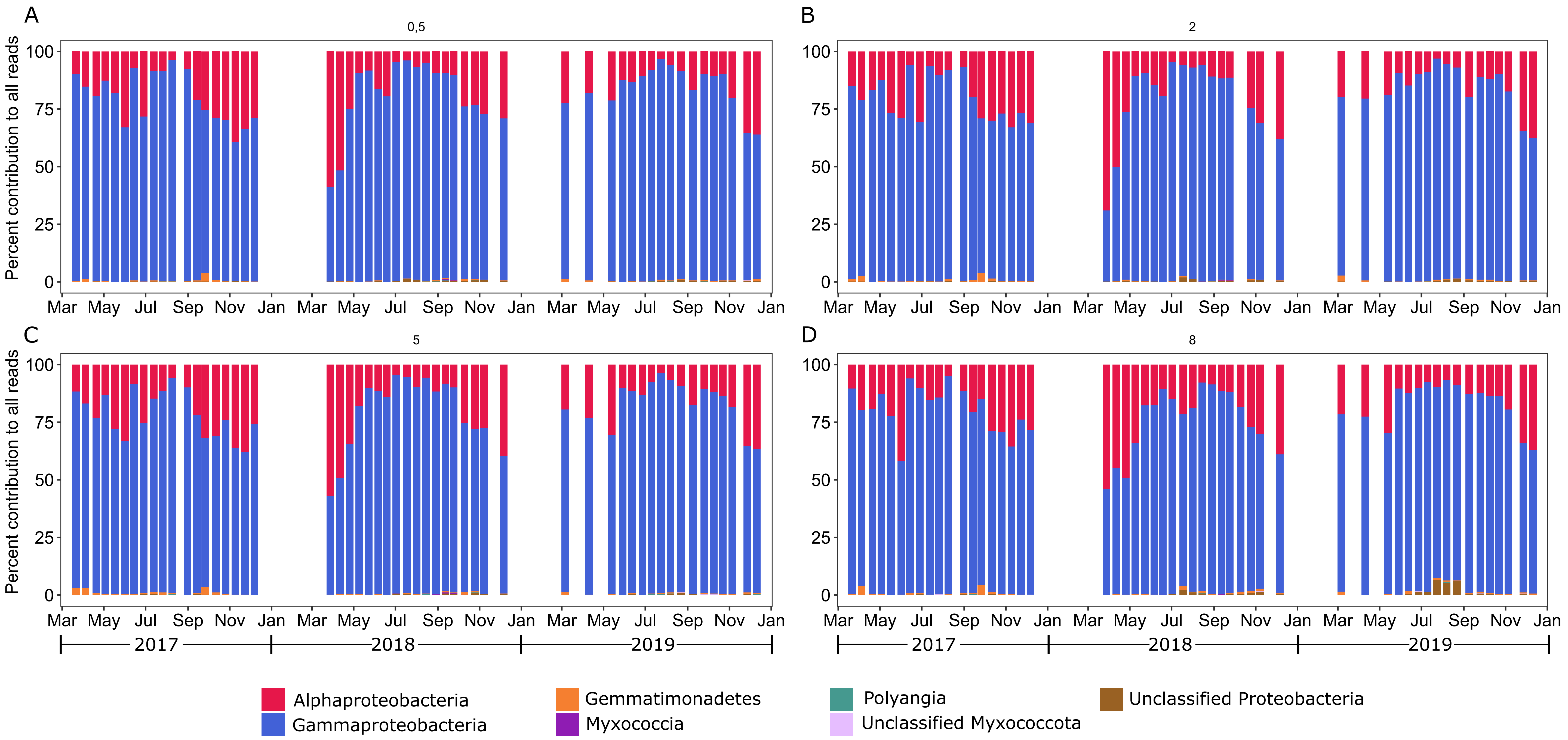


Supplementary Figure S8: AAP bacteria community composition according to *puf*M gene taxonomic assignment at class level for 0.5 (A), 2 (B), 5 (C) and 8 meters' depth (D) during 3-years sampling campaign.
