## Supplementary Figure S9 for "Phenology and ecological role of Aerobic Anoxygenic Phototrophs in fresh waters"

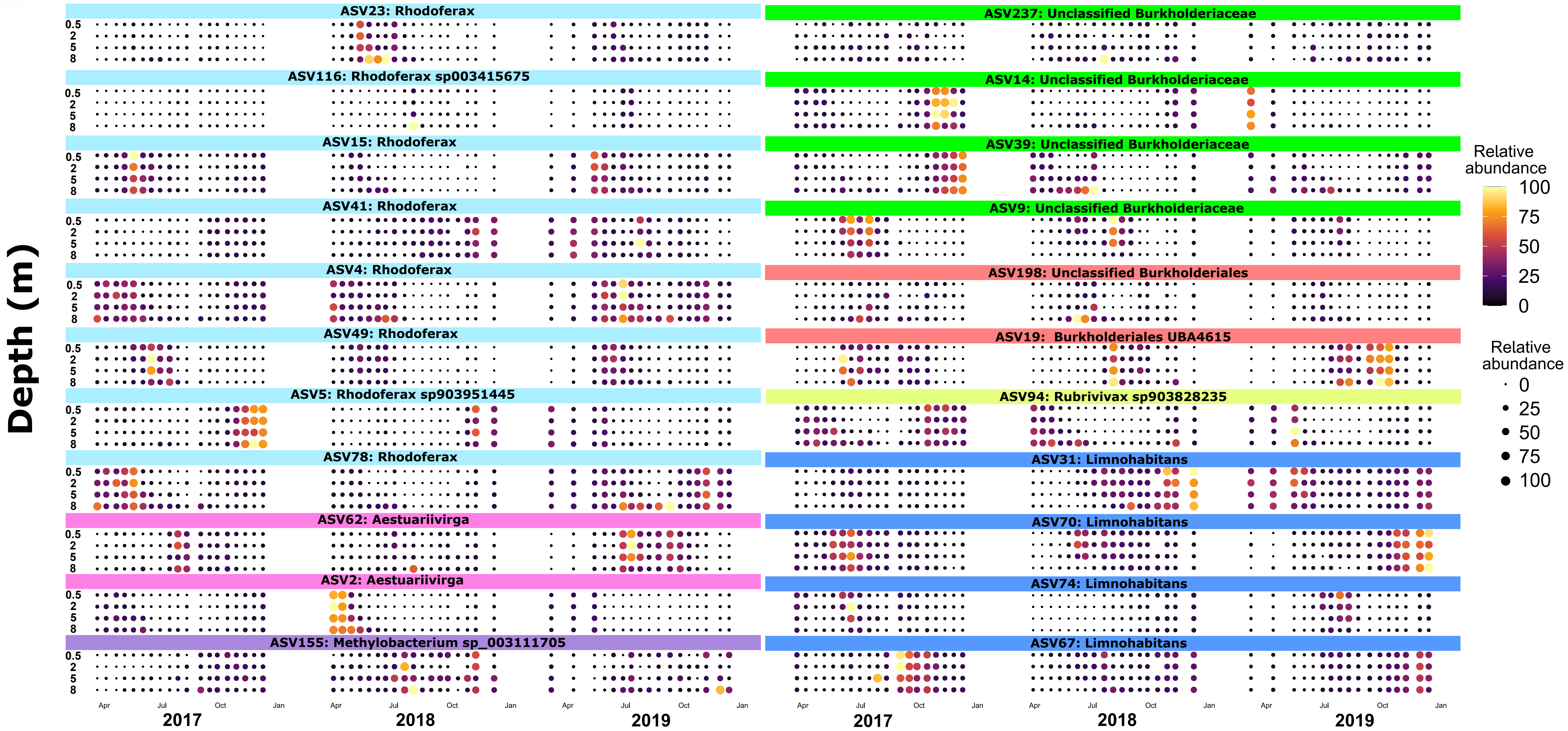


Supplementary Figure S9: Individually normalized relative abundance of the 22 core AAP ASVs during 3 years in 4 depths. Brighter colors and bigger dots indicate larger contribution to the AAP bacterial community. ASVs are clustered according to taxonomic classification at the maximum possible level (genus, family or order).
